# Technical Note: STRATIS: A Cloud-enabled Software Toolbox for Radiotherapy and Imaging Analysis

**DOI:** 10.1101/2022.11.08.515686

**Authors:** Aditya P. Apte, Eve LoCastro, Aditi Iyer, Jue Jiang, Jung Hun Oh, Harini Veeraraghavan, Amita Shukla-Dave, Joseph O. Deasy

## Abstract

**Purpose:** Recent advances in computational resources, including software libraries and hardware, have enabled the use of high-dimensional, multi-modal datasets to build Artificial Intelligence (AI) models and workflows for radiation therapy and image analysis. The purpose of Software Toolbox for RAdioTherapy and Imaging analysiS (STRATIS) is to provide cloud-enabled, easy-to-share software workflows to train and deploy AI models for transparency and multi-institutional collaboration.

**Method:** STRATIS leverages open source medical image informatics software for application-specific analysis. Jupyter notebooks for AI modeling workflows are provided with Python language as the base kernel. In addition to Python, workflows use software written in other languages, such as MATLAB, GNU-Octave, R, and C++, with the help of bridge libraries. The workflows can be run on a cloud platform, local workstation, or an institutional HPC cluster. Computational environments are provided in the form of publicly available docker images -and build scripts for local Anaconda environments. Utilities provided with STRATIS simplify bookkeeping of associations between imaging objects and allow chaining data processing operations defined via a setting file for AI models.

**Results:** Workflows available on STRATIS can be broadly categorized into image segmentation, deformable image registration, and outcomes modeling for radiotherapy toxicity and tumor control using radiomics and dosimetry features. The STRATIS-forge GitHub organization https://www.github.com/stratis-forge hosts build-scripts for Docker and Anaconda as well as Jupyter notebooks for analysis workflows. The software for building environments and workflow notebooks has open source-GNU-GPL copyright, and AI models retain the copyright chosen by their original developers.

**Conclusion:** STRATIS enables researchers to deploy and share AI modeling workflows for radiotherapy and image analysis. STRATIS is publicly available on Terra.bio’s FireCloud platform with a pre-deployed computational environment and on GitHub organization for users pursuing local deployment.

## Introduction

Rapid advances have been made in applying Artificial Intelligence (AI) to medical images. AI models improve clinical efficiency and, ultimately patient care by rapidly automating components of treatment planning. Automating the segmentation of organs at risk (OARs) and tumors, image registration, and prediction of normal tissue complication probability (NTCP) and tumor control probability (TCP) helps with their consistent, repeatable application to large datasets from multiple institutions. STRATIS fills the need for a software resource providing end-to-end workflows to facilitate the adoption and application of AI models on a cloud platform and locally. Many libraries are available to train AI models, such as PyTorch^1^, Tensorflow^2^, and Scikit-learn^3^, as well as medical imaging-specific software for visualization and processing. Open-source software tools such as 3DSlicer^4^, XNAT^5^, Computational Environment for Radiological Research (CERR)^6^, and Plastimatch^7^ are used for data management, processing, and visualization. MONAI^8^ is an open-source platform to train and deploy PyTorch-based convolutional neural networks (CNN) models, which offers step-by-step processing and training of medical images. The Insight ToolKit^9^ (ITK; www.itk.org) is an open-source, BSD-copyrighted software for image processing, with wrappers in commonly used, interpreted, and compiled languages. DeepInfer^10^ that allows users to utilize AI models from 3DSlicer and ModelHub^11^ can be used for web-based distribution of inference models.

Computational resources required for developing AI models are typically on the scale of institutional HPC or public cloud services such as Google Cloud Platform, Azure, or Amazon Web Services (AWS). The Imaging Data Commons (IDC)^12^ provides means to access data stored in public repositories such as The Cancer Imaging Archive (TCIA)^13^ from a cloud platform and provides examples of consuming and using the datasets for model development. Image data for training a model goes through various preprocessing transformations such as resampling, resizing, filtering, normalizing, harmonizing images, and combining multiple modalities, radiotherapy planning data, or time points. STRATIS simplifies data preparation for training an AI model and consumes its results by providing utilities for bookkeeping and image transformations. STRATIS workflows include all the steps required for pulling data from sources, training or inferring from an AI model, utilities for data management, visualization, and simplified access to image metadata.

### Building blocks of STRATIS workflow

STRATIS analysis workflows use open-source AI and image informatics libraries in a modular fashion. Figure 1 shows a schematic diagram of the building blocks. They broadly involve transforming the archived data, such as DICOM, to an analyzable format for preprocessing, model building or inference, and post-processing and visualizing results. The workflows are not tied to any specific processing or model building librar anduse fit-for-purpose software. CERR (cited 966 times as of Nov 2022) is used extensively for data management and transformations as it is well-tested and widely used for radiotherapy research. CERR’s GitHub webpage (https://github.com/cerr/CERR) was accessed 15463 times from 81 different countries over the last six months (Apr 27, 2022 – Oct 27, 2022) and cloned 372 times, excluding the zip file downloads and API calls. The workflows also use ITK for image input/output (I/O) and filtering, Plastimatch and Mermaid^14 15^ for image registration, and PyTorch and Tensorflow for model training. Results can be rendered directly in Jupyter notebooks and medical-specific viewers such as OHIF or 3DSlicer for review. As described in a later section, STRATIS provides utilities to pass imaging objects between various processing and visualization libraries. The workflows can use open datasets from TCIA via IDC utilities and XNAT via the REST API to pull and push data.

**Figure 1:**
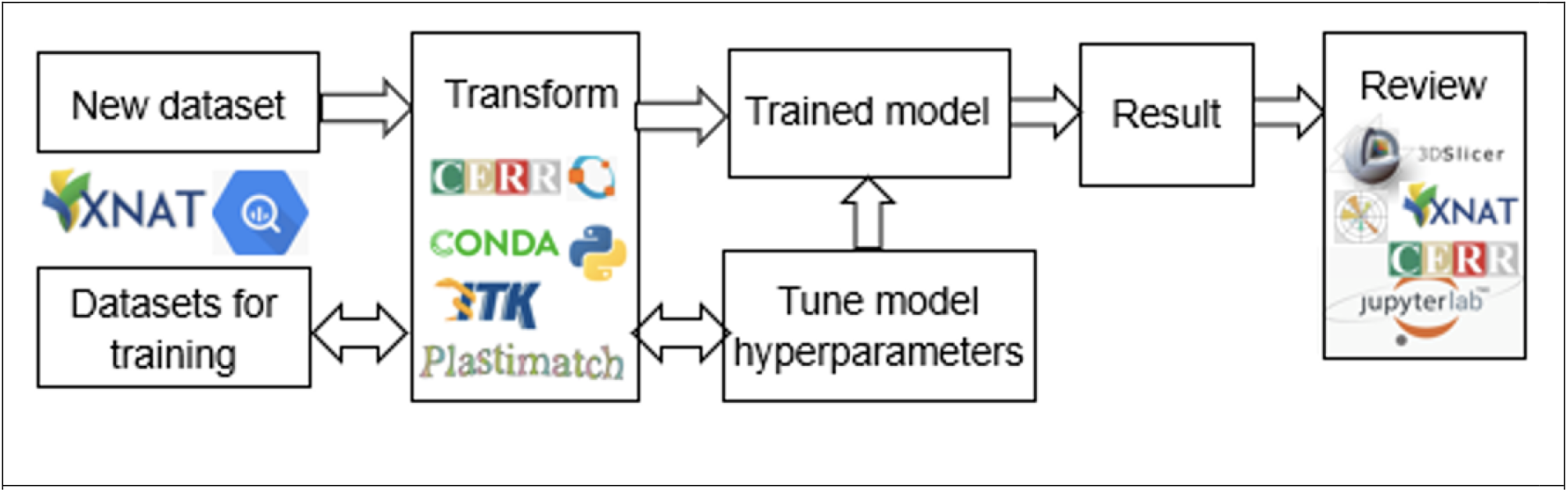
STRATIS workflows consist of modules for data transformations, model training, and deployment.

CERR software platform was originally developed in the MATLAB programming language and recently extended to be fully compatible with GNU-Octave. CERR users have the flexibility of invoking routines from Python using the Oct2py bridge library and Jupyter notebooks without a MATLAB license. CERR’s data structure is well suited to keep track of linkages between longitudinal and multi-modal images and radiation therapy planning data as it parses DICOM metadata. Automated pipelines, driven by JSON-format settings, were created in CERR to process radiotherapy data required for model building and to post-process results to associate with the desired frame of reference. CERR provides tools to use private vendor-specific DICOM metadata in MR images, which is necessary for derivation of diffusion and pharmacokinetic parametric maps from Diffusion Weighted and Dynamic Contrast-Enhanced MRI. CERR’s IBSI^16^ compatible radiomics and texture calculation are widely used^17^ (96 citations as of Nov 2022). In addition to texture maps, CERR provides tools for SUV computation from PET images. A library of image segmentation and outcomes model implementations^18^ was recently added to CERR.

FireCloud^19^ is an NCI Cloud Resource project powered by Terra^20^ for biomedical researchers to access data, run analysis tools, and collaborate. FireCloud provides computational environments for running Docker images containing analysis libraries and dependencies. Computational environments designed for STRATIS workflows are distributed publicly on the DockerHub platform (https://hub.docker.com). Each container can contain multiple versions of CUDA to allow different versions of modeling libraries.

FireCloud provides the means to seamlessly instantiate and run these images on the Google Cloud Platform (GCP) (https://cloud.google.com). The Workspaces on FireCloud containing STRATIS workflows can be shared between collaborators. Workflows can also be implemented to run locally via an Anaconda environment. Figure 2 shows the schematic of using STRATIS. Users can share data and Anaconda environments containing AI models within the workspace bucket. Additionally, users have the option to import datasets hosted in private data buckets on GCP and from other systems such as XNAT and IDC.

**Figure 2:**
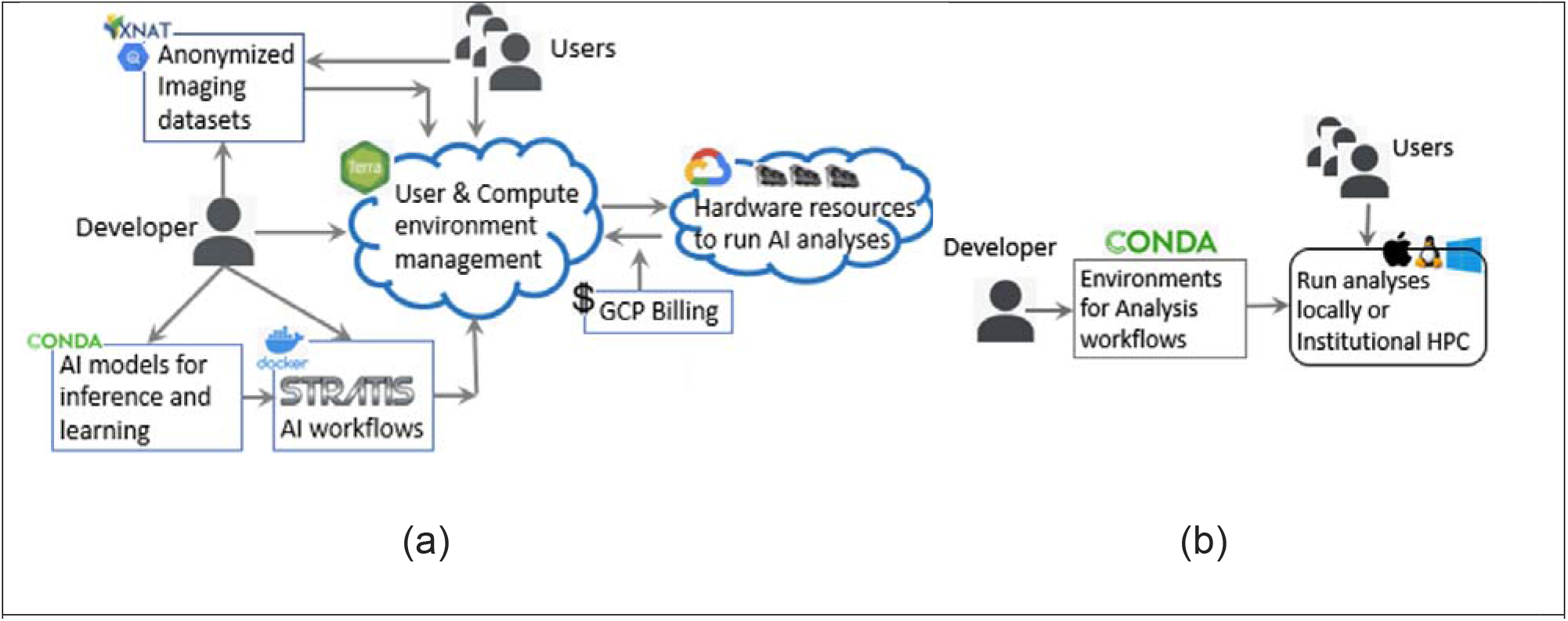
(a) STRATIS workflows are deployed to the FireCloud platform for instant use on user datasets. (b) STRATIS computational environments can be deployed locally on a workstation or institutional HPC resources.

#### STRASTIS-forge

The STRATIS-forge organization on GitHub hosts Jupyter notebooks^21^ containing the workflows and build-scripts requirements for Anaconda environments. These tools enable users to create local computational environments without FireCloud resources. A collection of analysis-specific repositories for image segmentation, registration, radiomics, and radiotherapy-dose features, radiotherapy normal tissue complication and tumor control estimation, data I/O through XNAT, and the interface between 3DSlicer and CERR for analysis and visualization are provided. Table 1 provides the list of available models in STRATIS.

**Table 1:**
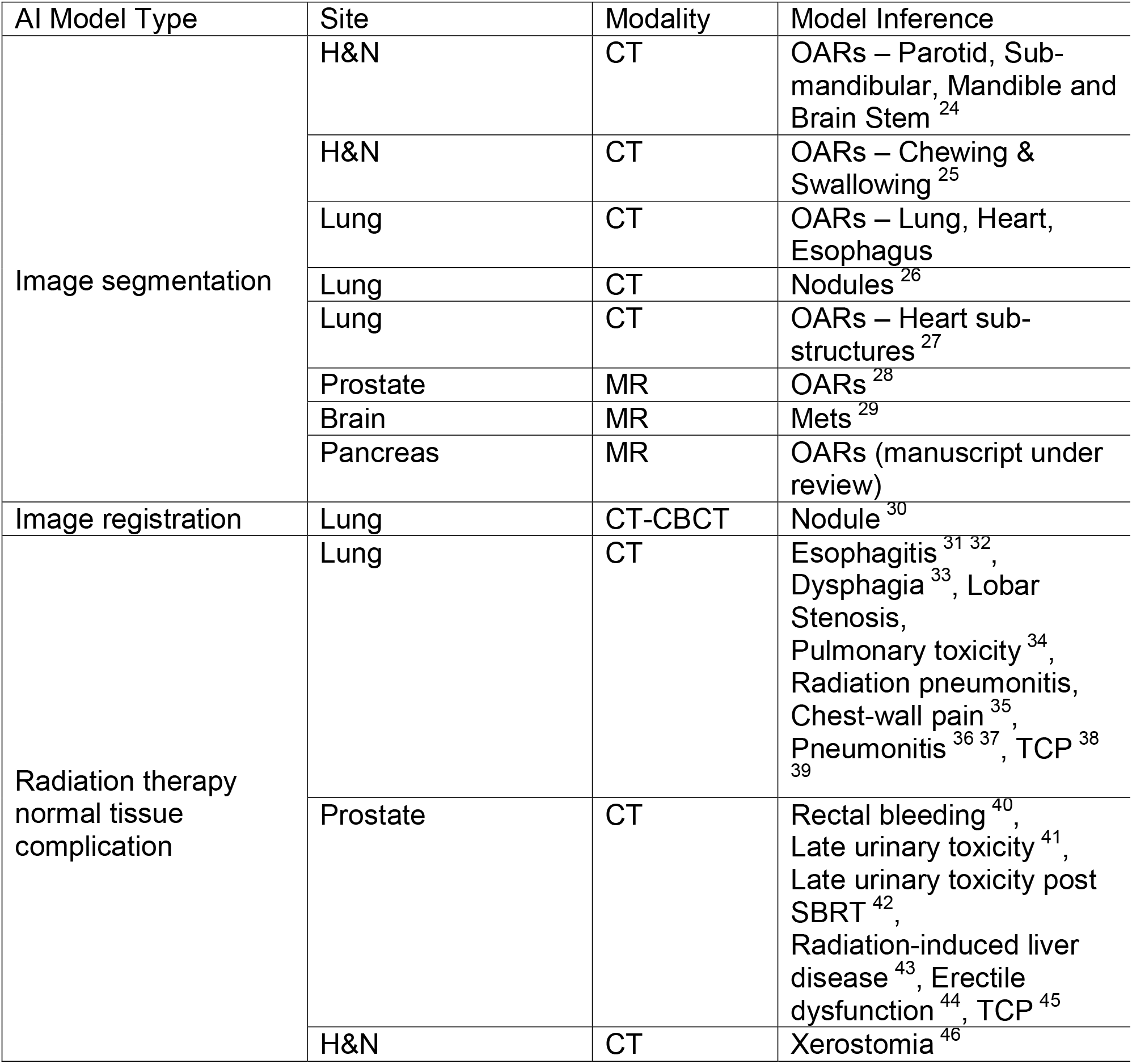
AI models available in STRATIS.

Image segmentation models in STRATIS are available for multiple modalities such as CT, MR, and CBCT and for different sites, including Head and Neck (H&N), Lung, Prostate, Brain, and Pancreas. Image registration models are available for longitudinal and CT-CBCT applications. Figure 3 shows an output of the CT-CBCT registration model to a longitudinal patient dataset from our institution. The magnitude of deformation increases as treatment progresses, capturing the shrinkage of the tumor. A STRATIS workflow is provided, which can be applied to other user datasets and publicly available longitudinal lung imaging datasets such as 4DLung^22^ collection from TCIA. Many of the image segmentation models in STRATIS have been used clinically at our institution in radiotherapy planning workflows to automate OAR segmentation for planner review.

**Figure 3:**
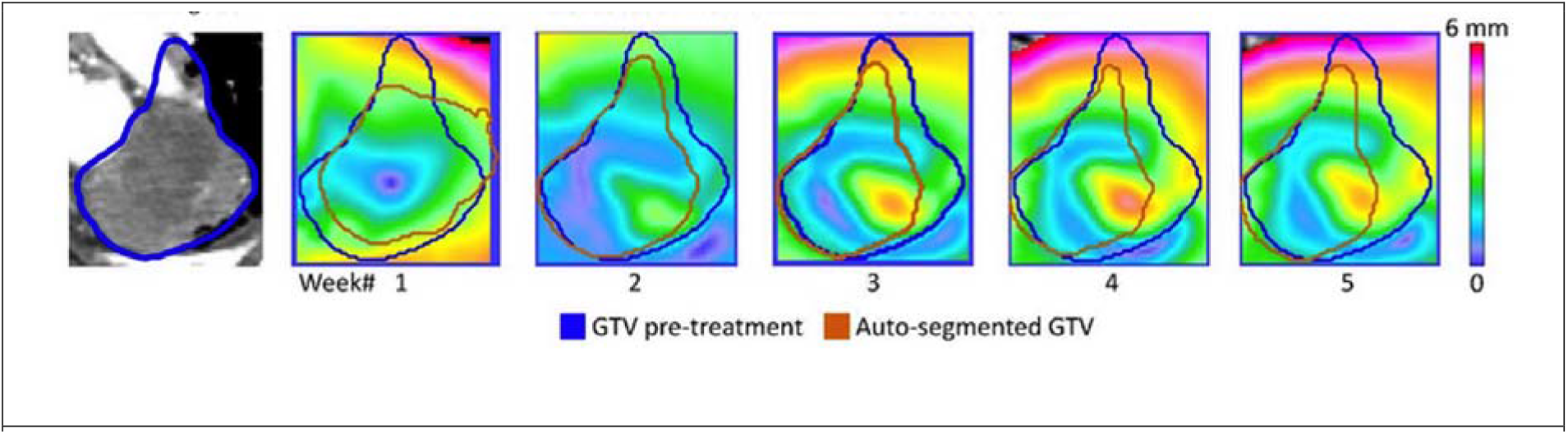
Magnitude of deformation vector field derived from applying the CT-CBCT image registration model to CBCTs acquired over 5 weeks from treatment planning.

Some models have also been validated on datasets from other institutions^23^ demonstrating their robustness and reproducibility. NTCP and TCP models are based on radiotherapy dose and radiomics features. Workflows allow users to investigate the impact of scaling dose on predicted NTCP and TCP, allowing researchers to personalize radiotherapy treatment prescriptions.

Figure 4 shows sample dose-response curves for grade-2 acute esophagitis and radiation pneumonitis, along with predicted TCP in a patient with stage-III non-small cell lung cancer (NSCLC).

**Figure 4:**
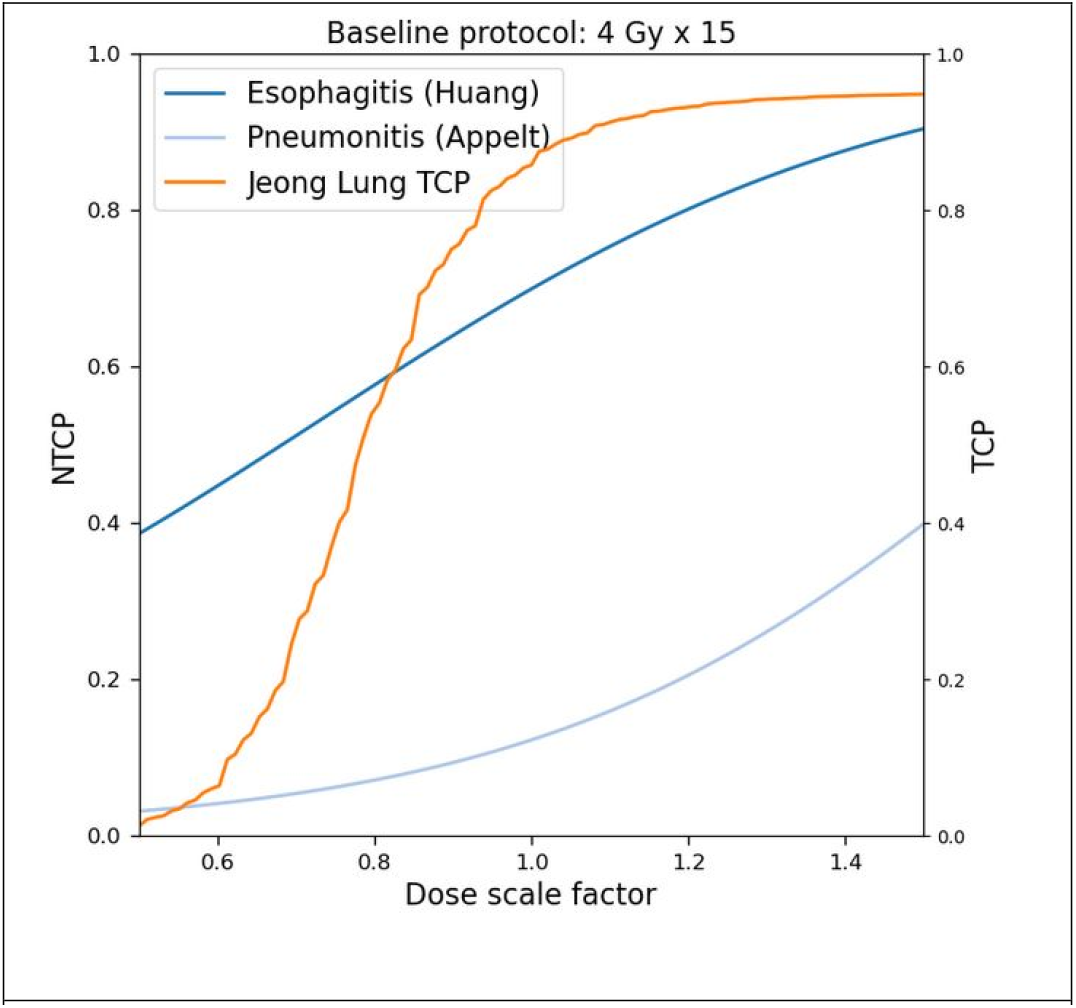
Dose-response for a stage-III NSCLC patient to simulate the effect of scaling fractional dose on NTCP and TCP.

#### Interfacing with open-source image informatics software

STRATIS provides tools to exchange data types between CERR and 3DSlicer and archival and use of images from the XNAT platform. The next sections describe these utilities in more detail.

##### CERR-3DSlicer data exchange

3DSlicer is a widely used software for the visualization of medical images with a vast number of extensions for processing and analysis. We developed *sliCERR*, an extension written in Python to facilitate access to CERR’s radiotherapy and image analysis functionality and to review the computational results from CERR in 3DSlicer. *sliCERR* uses *oct2py* and octave_kernel packages in the Slicer Python interpreter via the *pip_install* feature and GNU-Octave and CERR. The *sliCERR* extension is platform-agnostic and works on Windows, MacOS or Linux. *sliCERR* provides various modules to facilitate data I/O conversion. The cerr2mrml module handles the I/O of loading native CERR planC format files into the 3D Slicer MRML scene, including import of scan, radiotherapy-dose and region of interest (ROI) contours. Use of recent developments in CERR such as Deep Learning-based image segmentation and radiomics texture mapping, the Radiotherapy Outcomes Estimator (ROE) and semi-quantitative DCE features is demonstrated in Jupyter notebooks publicly available on STRATIS-forge. Figure 5 shows an example of visualizing scan, radiotherapy-dose and structures from CERR’s planC object in 3DSlicer. The sliCERR extension is available for preview on GitHub (http://github.com/cerr/sliCERR) and upon release, will be directly available from the Slicer Extension Manager. https://github.com/cerr/slicerr/blob/main/slicer_io_example.py notebook demonstrates the use of sliCERR to import CERR metadata in 3DSlicer.

**Figure 5:**
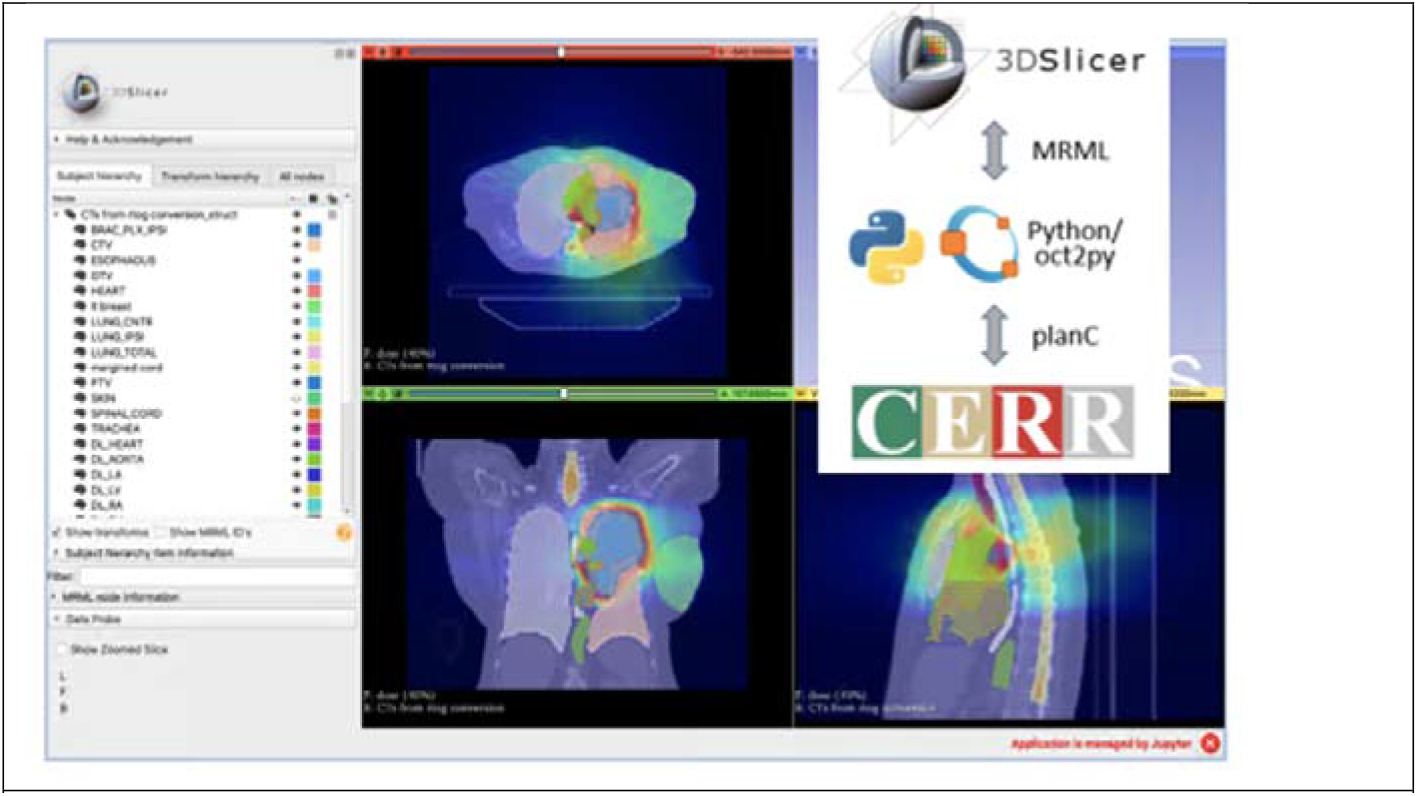
An example of importing metadata from CERR’s planC data structure in 3DSlicer as MRML scene.

##### XNAT usage in STARTIS

XNAT is a web-based platform for medical image management and collaboration. We host an instance of XNAT v1.8.1, christened the “Predictive Informatics XNAT” (“PIXNAT”) for collaborative medical physics research. The PIXNAT is institutionally approved for housing research imaging data and accessible to external researchers for data sharing and processing. All data is fully anonymized using Clinical Trials Processor (CTP)^47^ according to institutional guidelines. PIXNAT is accessible at http://pixnat.mskcc.org. Currently, it houses 42 projects, 3366 subjects, and 7405 imaging sessions. It has been used to collaborate with ten different institutions.

Analysis-ready data from PIXNAT can be used in STRATIS workflows. Similarly, results from STRATIS can be pushed to PIXNAT for archival. PIXNAT is extended with OHIF-XNAT Viewer and ROI datatype plugins to facilitate the overlay of segmentation on images. Figure 6 shows the overlay of segmentation results from the cardiac sub-structure model in the OHIF-XNAT viewer. https://github.com/stratis-forge/xnat-workflows/blob/main/xnat_data_pull_demo.ipynb shows the use of images from PIXNAT to interactively run Python and CERR routines and visualize in-line results.

**Figure 6:**
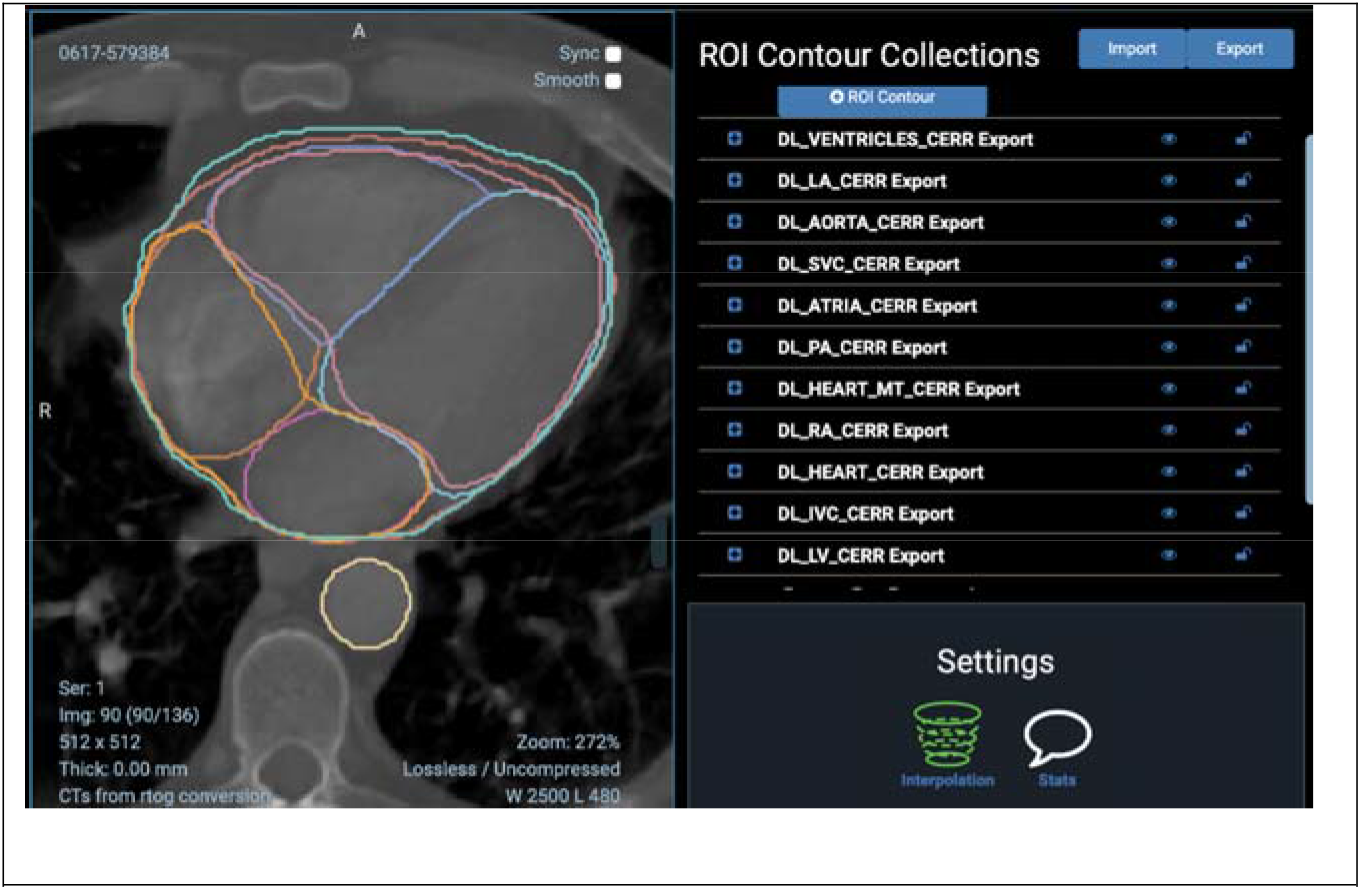
An example of displaying AI generated segmentation of cardiac sub-structures in XNAT-OHIF viewer.

### Conclusion

STRATIS provides readily usable and reproducible AI workflows for radiotherapy and image analysis. Use cases with examples were presented to demonstrate the use of STRATIS on publicly available Terra.bio’s FireCloud platform as well as local deployment. STRATIS will ultimately help to speed-up the adoption of AI models by making complete analysis pipelines easily accessible to researchers.

## Acknowledgments

This research was partially funded by NIH/NCI Cancer Center Support grant P30 CA008748.

## References

1 Adam Paszke, Sam Gross, Francisco Massa, Adam Lerer, James Bradbury, Gregory Chanan, Trevor Killeen, Zeming Lin, Natalia Gimelshein, Luca Antiga, Alban Desmaison, Andreas Köpf, Edward Yang, Zach DeVito, Martin Raison, Alykhan Tejani, Sasank Chilamkurthy, Benoit Steiner, Lu Fang, Junjie Bai, and Soumith Chintala, “PyTorch: An Imperative Style, High-Performance Deep Learning Library”, (2019).

2 Martin and Barham Abadi, Paul and Chen, Jianmin and Chen, Zhifeng and Davis, Andy and Dean, Jeffrey and Devin, Matthieu and Ghemawat, Sanjay and Irving, Geoffrey and Isard, Michael and others, “Tensorflow: A system for large-scale machine learning”, in Symposium on Operating Systems Design and Implementation (2016), pp. 265–283.

3 F. Pedregosa, G. Varoquaux, A. Gramfort, V. Michel, B. Thirion, O. Grisel, M. Blondel, P. Prettenhofer, R. Weiss, V. Dubourg, J. Vanderplas, A. Passos, D. Cournapeau, M. Brucher, M. Perrot, and E. Duchesnay, “Scikit-learn: Machine Learning in Python,” J Mach Learn Res 12, 2825–2830 (2011).

4 Pieper SD Kikinis R, Vosburgh K, “3D Slicer: A Platform for Subject-Specific Image Analysis, Visualization, and Clinical Support,” In: Jolesz F. (eds) Intraoperative Imaging and Image-Guided Therapy (2014).

5 D. S. Marcus, T. R. Olsen, M. Ramaratnam, and R. L. Buckner, “The Extensible Neuroimaging Archive Toolkit - An informatics platform for managing, exploring, and sharing neuroimaging data,” Neuroinformatics 5 (1), 11–33 (2007).

6 J. O. Deasy, A. I. Blanco, and V. H. Clark, “CERR: a computational environment for radiotherapy research,” Med Phys 30 (5), 979–985 (2003).

7 P. Zaffino, P. Raudaschl, K. Fritscher, G. C. Sharp, and M. F. Spadea, “Technical Note: plastimatch mabs, an open source tool for automatic image segmentation,” Med Phys 43 (9), 51–55 (2016).

8 Nic Ma; Wenqi Li; Richard Brown; Yiheng Wang; Benjamin Gorman; Behrooz; Hans Johnson; Isaac Yang; Eric Kerfoot; Yiwen Li; Mohammad Adil; Yuan-Ting Hsieh (Wł5û5ĩ); charliebudd; Arpit Aggarwal; Cameron Trentz; adam aji; Ben Murray; Gagan Daroach; Petru-Daniel Tudosiu; myron; Mark Graham; Balamurali; Christian Baker; Jan Sellner; Lucas Fidon; Alex Powers; Guy Leroy; Alxaline; Daniel Schulz;, 2020.

9 W. Schroeder L. Ibáñez, L. Ng, and J. Cates, 2003.

10 A. Mehrtash, M. Pesteie, J. Hetherington, P. A. Behringer, T. Kapur, W. M. Wells, 3rd, R. Rohling, A. Fedorov, and P. Abolmaesumi, “DeepInfer: Open-Source Deep Learning Deployment Toolkit for Image-Guided Therapy,” Proc SPIE Int Soc Opt Eng 10135 (2017).

11 Michael Schwier Ahmed Hosny, Christoph Berger, Evin P Örnek, Mehmet Turan, et al, “ModelHub.AI: Dissemination Platform for Deep Learning Models,” arXiv preprint arXiv:l9ll.13218 (2019).

12 A. Fedorov, W. Longabaugh, D. Pot, D. Clunie, S. Pieper, R. Lewis, H. Aerts, A. Homeyer, M. Herrmann, U. Wagner, T. Pihl, K. Farahani, and R. Kikinis, “NCI Imaging Data Commons,” International Journal of Radiation Oncology Biology Physics 111 (3), E101–E101 (2021).

13 K. Clark, B. Vendt, K. Smith, J. Freymann, J. Kirby, P. Koppel, S. Moore, S. Phillips, D. Maffitt, M. Pringle, L. Tarbox, and F. Prior, “The Cancer Imaging Archive (TCIA): Maintaining and Operating a Public Information Repository,” J Digit Imaging 26 (6), 1045–1057 (2013).

14 X. Han, J. Hong, M. Reyngold, C. Crane, J. Cuaron, C. Hajj, J. Mann, M. Zinovoy, H. Greer, E. Yorke, G. Mageras, and M. Niethammer, “Deep-learning-based image registration and automatic segmentation of organs-at-risk in cone-beam CT scans from high-dose radiation treatment of pancreatic cancer,” Medical Physics 48 (6), 3084–3095 (2021).

15 Z. Y. Shen, X. Han, Z. L. Xu, and M. Niethammer, “Networks for Joint Affine and Non-parametric Image Registration,” Proc Cvpr leee, 4219–4228 (2019).

16 A. Zwanenburg, M. Vallieres, M. A. Abdalah, H. J. W. L. Aerts, V. Andrearczyk, A. Apte, S. Ashrafinia, S. Bakas, R. Beukinga, R. Boellaard, M. Bogowicz, L. Boldrini, I. Buvat, G. J. R. Cook, C. Davatzikos, A. Depeursinge, M. C. Desseroit, N. Dinapoli, C. V. Dinh, S. Echegaray, I. El Naqa, A. Y. Fedorov, R. Gatta, R. J. Gillies, V. Goh, M. Gotz, M. Guckenberger, S. M. Ha, M. Hatt, F. Isensee, P. Lambin, S. Leger, R. T. Leijenaar, J. Lenkowicz, F. Lippert, A. Losnegard, K. H. Maier-Hein, O. Morin, H. Muller, S. Napel, C. Nioche, F. Orlhac, S. Pati, E. A. Pfaehler, A. Rahmim, A. U. Rao, J. Scherer, M. M. Siddique, N. M. Sijtsema, J. S. Fernandez, E. Spezi, R. J. Steenbakkers, S. Tanadini-Lang, D. Thorwarth, E. G. Troost, T. Upadhaya, V. Valentini, L. V. van Dijk, J. van Griethuysen, F. H. van Velden, P. Whybra, C. Richter, and S. Lock, “The Image Biomarker Standardization Initiative: Standardized Quantitative Radiomics for High-Throughput Image-based Phenotyping,” Radiology 295 (2), 328–338 (2020).

17 A. P. Apte, A. Iyer, M. Crispin-Ortuzar, R. Pandya, L. V. van Dijk, E. Spezi, M. Thor, H. Um, H. Veeraraghavan, J. H. Oh, A. Shukla-Dave, and J. O. Deasy, “Technical Note: Extension of CERR for computational radiomics: A comprehensive MATLAB platform for reproducible radiomics research,” Med Phys (2018).

18 A. P. Apte, A. Iyer, M. Thor, R. Pandya, R. Haq, J. Jiang, E. LoCastro, A. Shukla-Dave, N. Sasankan, Y. Xiao, Y. C. Hu, S. Elguindi, H. Veeraraghavan, J. H. Oh, A. Jackson, and J. O. Deasy, “Library of deep-learning image segmentation and outcomes model-implementations,” Phys Medica 73, 190–196 (2020).

19 Megan Hanna Chet Birger, Edward Salinas, Jason Neff, Gordon Saksena, Dimitri Livitz, Daniel Rosebrock, View ORCID ProfileChip Stewart, Ignaty Leshchiner, Alexander Baumann, Douglas Voet, Kristian Cibulskis, Eric Banks, Anthony Philippakis, Gad Getz, “FireCloud, a scalable cloud-based platform for collaborative genome analysis: Strategies for reducing and controlling costs,” bioRxiv (2017).

20 Jeffrey M. Perkel, “Terra takes the pain out of ‘omics’ computing in the cloud,” Nature 601, 154–155 (2022).

21 T. Kluyver, B. Ragan-Kelley, F. Perez, B. Granger, M. Bussonnier, J. Frederic, K. Kelley, J. Hamrick, J. Grout, S. Corlay, P. Ivanov, D. Avila, S. Abdalla, C. Willing, and Jupyter Dev Team, “Jupyter Notebooks-a publishing format for reproducible computational workflows,” Positioning and Power in Academic Publishing: Players, Agents and Agendas, 87–90 (2016).

22 G. D. Hugo, S. Balik, P. J. Keall, J. Lu, and J. F. Williamson, “A longitudinal four-dimensional computed tomography and cone beam computed tomography dataset for image-guided radiation therapy research in lung cancer,” Medical Physics 44 (2), 762–771 (2017).

23 G. M. Walls, V. Giacometti, A. Apte, M. Thor, C. McCann, G. G. Hanna, J. ‘Connor, J. O. Deasy, A. R. Hounsell, K. T. Butterworth, A. J. Cole, S. Jain, and C. K. McGarry, “Validation of an established deep learning auto-segmentation tool for cardiac substructures in 4D radiotherapy planning scans,” Phys Imag Radiat Onc 23, 118–126 (2022).

24 J. Jiang, S. Elguindi, S. L. Berry, I. Onochie, L. Cervino, J. O. Deasy, and H. Veeraraghavan, “Nested block self-attention multiple resolution residual network for multiorgan segmentation from CT,” Medical Physics 49 (8), 5244–5257 (2022).

25 A. Iyer, M. Thor, I. Onochie, J. Hesse, K. Zakeri, E. LoCastro, J. Jiang, H. Veeraraghavan, S. Elguindi, N. Y. Lee, J. O. Deasy, and A. P. Apte, “Prospectively-validated deep learning model for segmenting swallowing and chewing structures in CT,” Phys Med Biol 67 (2) (2022).

26 J. Jiang, Y. C. Hu, C. J. Liu, D. Halpenny, M. D. Hellmann, J. O. Deasy, G. Mageras, and H. Veeraraghavan, “Multiple Resolution Residually Connected Feature Streams for Automatic Lung Tumor Segmentation From CT Images,” leee T Med Imaging 38 (1), 134–144 (2019).

27 R. Haq, A. Hotca, A. Apte, A. Rimner, J. O. Deasy, and M. Thor, “Cardio-pulmonary substructure segmentation of radiotherapy computed tomography images using convolutional neural networks for clinical outcomes analysis,” Phys Imaging Radiat Oncol 14, 61–66 (2020).

28 S. Elguindi, M. J. Zelefsky, J. Jiang, H. Veeraraghavan, J. O. Deasy, M. A. Hunt, and N. Tyagi, “Deep learning-based auto-segmentation of targets and organs-at-risk for magnetic resonance imaging only planning of prostate radiotherapy,” Phys Imag Radiat Onc 12, 80–86 (2019).

29 D. G. Hsu, A. Ballangrud, A. Shamseddine, J. O. Deasy, H. Veeraraghavan, L. Cervino, K. Beal, and M. Aristophanous, “Automatic segmentation of brain metastases using T1 magnetic resonance and computed tomography images,” Phys Med Biol 66 (17) (2021).

30 J. Jiang and H. Veeraraghavan, “One Shot PACS: Patient Specific Anatomic Context and Shape Prior Aware Recurrent Registration-Segmentation of Longitudinal Thoracic Cone Beam CTs,” leee T Med Imaging 41 (8), 2021–2032 (2022).

31 E. X. Huang, A. J. Hope, P. E. Lindsay, M. Trovo, I. El Naqa, J. O. Deasy, and J. D. Bradley, “Heart irradiation as a risk factor for radiation pneumonitis,” Acta Oncologica 50 (1), 51–60 (2011).

32 R. Wijsman, F. Dankers, E. G. C. Troost, A. L. Hoffmann, E. H. F. M. van der Heijden, L. F. de Geus-Oei, and J. Bussink, “Multivariable normal-tissue complication modeling of acute esophageal toxicity in advanced stage non-small cell lung cancer patients treated with intensity-modulated (chemo-)radiotherapy,” Radiotherapy and Oncology 117 (1), 49–54 (2015).

33 T. Rancati, M. Schwarz, A. M. Allen, F. Feng, A. Popovtzer, B. Mittal, and A. Eisbruch, “Radiation Dose-Volume Effects in the Larynx and Pharynx,” International Journal of Radiation Oncology Biology Physics 76 (3), S64–S69 (2010).

34 H. Tekatli, M. Duijm, E. O. D. Hoop, W. Verbakel, W. Schillemans, B. J. Slotman, J. J. Nuyttens, and S. Senan, “Normal Tissue Complication Probability Modeling of Pulmonary Toxicity After Stereotactic and Hypofractionated Radiation Therapy for Central Lung Tumors,” International Journal of Radiation Oncology Biology Physics 100 (3), 738–747 (2018).

35 S. U. Din, E. L. Williams, A. Jackson, K. E. Rosenzweig, A. J. Wu, A. Foster, E. D. Yorke, and A. Rimner, “Impact of Fractionation and Dose in a Multivariate Model for Radiation-Induced Chest Wall Pain,” International Journal of Radiation Oncology Biology Physics 93 (2), 418–424 (2015).

36 A. L. Appelt, I. R. Vogelius, K. P. Farr, A. A. Khalil, and S. M. Bentzen, “Towards individualized dose constraints: Adjusting the QUANTEC radiation pneumonitis model for clinical risk factors,” Acta Oncologica 53 (5), 605–612 (2014).

37 M. Thor, J. Deasy, A. Iyer, E. Bendau, A. Fontanella, A. Apte, E. Yorke, A. Rimner, and A. Jackson, “Toward personalized dose-prescription in locally advanced non-small cell lung cancer: Validation of published normal tissue complication probability models,” Radiotherapy and Oncology 138, 45–51 (2019).

38 A. Fontanella, C. Robinson, A. Zuniga, A. Apte, W. Thorstad, J. Bradley, and J. Deasy, “Test of the Generalized Tumor Dose (gTD) Model with An Independent Lung Tumor Dataset,” Medical Physics 41 (6), 296–+ (2014).

39 J. Jeong, J. H. Oh, J. J. Sonke, J. Belderbos, J. D. Bradley, A. N. Fontanella, S. S. Rao, and J. O. Deasy, “Modeling the Cellular Response of Lung Cancer to Radiation Therapy for a Broad Range of Fractionation Schedules,” Clinical Cancer Research 23 (18), 5469–5479 (2017).

40 J. M. Michalski, H. Gay, A. Jackson, S. L. Tucker, and J. O. Deasy, “Radiation Dose-Volume Effects in Radiation-Induced Rectal Injury,” International Journal of Radiation Oncology Biology Physics 76 (3), S123–S129 (2010).

41 M. R. Cheung, S. L. Tucker, L. Dong, R. de Crevoisier, A. K. Lee, S. Frank, R. J. Kudchadker, H. Thames, R. Mohan, and D. Kuban, “Investigation of bladder dose and volume factors influencing late urinary toxicity after external beam radiotherapy for prostate cancer,” International Journal of Radiation Oncology Biology Physics 67 (4), 1059–1065 (2007).

42 T. P. Kole, M. Tong, B. B. Wu, S. Y. Lei, O. Obayomi-Davies, L. N. Chen, S. M. Suy, A. Dritschilo, E. Yorke, and S. P. Collins, “Late urinary toxicity modeling after stereotactic body radiotherapy (SBRT) in the definitive treatment of localized prostate cancer,” Acta Oncologica 55 (1), 52–58 (2016).

43 C. C. Pan, B. D. Kavanagh, L. A. Dawson, X. A. Li, S. K. Das, M. Miften, and R. K. Ten Haken, “Radiation-Associated Liver Injury,” International Journal of Radiation Oncology Biology Physics 76(3), S94–S100 (2010).

44 M. Roach, D. M. Chinn, J. Holland, and M. Clarke, “A pilot survey of sexual function and quality of life following 3D conformal radiotherapy for clinically localized prostate cancer,” International Journal of Radiation Oncology Biology Physics 35 (5), 869–874 (1996).

45 S. Walsh, E. Roelofs, P. Kuess, P. Lambin, B. Jones, D. Georg, and F. Verhaegen, “A validated tumor control probability model based on a meta-analysis of low, intermediate, and high-risk prostate cancer patients treated by photon, proton, or carbon-ion radiotherapy,” Medical Physics 43 (2), 734–747 (2016).

46 K. S. C. Chao, J. O. Deasy, J. Markman, J. Haynie, C. A. Perez, J. A. Purdy, and D. A. Low, “A prospective study of salivary function sparing in patients with head-and-neck cancers receiving intensity-modulated or three-dimensional radiation therapy: Initial results,” International Journal of Radiation Oncology Biology Physics 49 (4), 907–916 (2001).

47 J. B. Freymann, J. S. Kirby, J. H. Perry, D. A. Clunie, and C. C. Jaffe, “Image data sharing for biomedical research--meeting HIPAA requirements for De-identification,” J Digit Imaging 25 (1), 14–24 (2012).

